# Kinetic modelling of quantitative proteome data predicts metabolic reprogramming of liver cancer

**DOI:** 10.1101/275040

**Authors:** Nikolaus Berndt, Antje Egners, Guido Mastrobuoni, Olga Vvedenskaya, Athanassios Fragoulis, Aurélien Dugourd, Sascha Bulik, Matthias Pietzke, Chris Bielow, Rob van Gassel, Steven Olde Damink, Merve Erdem, Julio Saez-Rodriguez, Hermann-Georg Holzhütter, Stefan Kempa, Thorsten Cramer

## Abstract

Metabolic alterations can serve as targets for diagnosis and therapy of cancer. Due to the highly complex regulation of cellular metabolism, definite identification of metabolic pathway alterations remains challenging and requires sophisticated experimentation. Here, we applied a comprehensive kinetic model of the central carbon metabolism (CCM) to characterize metabolic reprogramming in murine liver cancer. We show that relative differences of protein abundances of metabolic enzymes obtained by mass spectrometry can be used to scale maximal enzyme capacities. Model simulations predicted tumor - specific alterations of various components of the CCM, a selected number of which were subsequently verified by *in vitro* and *in vivo* experiments. Furthermore, we demonstrate the ability of the kinetic model to identify metabolic pathways whose inhibition results in selective tumor cell killing. Our systems biology approach establishes that combining cellular experimentation with computer simulations of physiology-based metabolic models enables a comprehensive understanding of deregulated energetics in cancer.

## Introduction

Worldwide, hepatocellular carcinoma (HCC) is the fifth most common cancer and the third most common cause of cancer - related deaths (Waller et al., 2015). Countless initiatives around the world in conjunction with the unprecedented development of high - throughput analytical methodology have spiraled up the molecular knowledge about cancer in dizzying heights (Garraway and Lander, 2013). Clinical translation of the newly gained information has resulted in the approval of a plethora of molecular - targeted drugs with antiproliferative activity. However, despite all these advances, the overall death rate for cancer has declined at a much slower pace in the past 40 years compared to other major causes of mortality such as cardiovascular and infectious diseases (Ma et al., 2015). This is - to a large extent – explained by suboptimal long - term antiproliferative efficacy of newly developed molecular - targeted drugs (Fojo and Parkinson, 2010). Cancer cells display a marked capability to compensate for the inactivation of signaling pathways - and other growth - promoting mechanisms - that are considered essential for neoplastic progression (McIntyre and Harris, 2015; Niewerth et al., 2015). This translates into the emergence of therapy resistance, a major obstacle of clinical oncology and a central hallmark of human HCC (Waller et al., 2015). To achieve effective and long - lasting therapy responses it is therefore of pivotal importance to identify processes that are at the same time essential and unique, thereby avoiding resistance via usage of alternative processes. Metabolism fulfills these characteristics and is indeed widely considered to represent an attractive target for cancer therapy (Schulze and Harris, 2012).

The notion that tumors display specific metabolic alterations that can be exploited for diagnosis and therapy of cancer has received widespread attention in recent years (Pavlova and Thompson, 2016). However, it also became evident that a reliable analysis of metabolism, especially under *in vivo* situations, is challenging to perform. This is due to various factors, e.g. the rapid turnover of substrates and products, the intricate complexity of the metabolic network, the central importance of external stimuli such as hormones, growth factors and the cellular microenvironment (Cazzaniga et al., 2014). One main reason for the insufficient understanding of metabolic changes in tumors is the strong focus on changes in the expression level of metabolic enzymes and transporters. The importance of downstream kinetic regulation, e.g. by allosteric effects or reversible phosphorylation, has been underestimated or completely disregarded in the last two decades (Weinberg, 2010). Choosing glucose metabolism of the liver as an example, we have recently demonstrated the necessity to combine existing knowledge on gene expression changes with the complex kinetic regulation of enzymes in order to understand the metabolic response of the liver to varying external challenges (Bulik et al., 2016). Here, we present an innovative concerted approach to study cancer metabolism by combining a novel physiology - based kinetic model of the central metabolism with high - quality quantitative proteomics data and molecular biological experimentation to elucidate metabolic differences between HCC and the normal liver in a murine HCC model. Using relative changes in the expression level of metabolic enzymes in HCC and normal liver cells to scale maximal enzyme activities, we simulate the metabolic response of HCC and the normal liver to variations in the metabolite and hormone profile of the blood plasma. This enables the definition of conditions at which the metabolism of the tumor becomes severely impaired while the metabolism of normal liver cells remains largely unaffected. We strongly believe that the herewith presented approach bears translational potential and will outline a basic roadmap to achieve this.

## Results

### Proteome analysis of the central carbon metabolism in normal and malignant murine liver

In order to generate a detailed expression profile of enzymes of the central carbon metabolism, HCC samples from ASV - B mice as well as liver tissue from tumor - free control mice were analyzed by a mass spectrometry - based shotgun proteomics approach (Cox and Mann, 2011). The proteomic experimental design consisted of 2 conditions (tumor and control), 7 biological replicates per condition and 2 technical replicates per biological replicate. In total, 4.415 proteins were detected across the 28 samples. Each pair of technical replicates was averaged resulting in 14 biological samples (Fig. S1A, S1B). Principal Component Analysis showed that the variance of the mouse samples was well explained by their status (control / tumor). The control and tumor samples are clearly separated along the first principal component, which explains 54% of the variance of the dataset (Fig. 1A, S1C). Furthermore, the second component shows that the tumor samples display a greater inter - sample variability than the control samples, as expected given the aberrant regulation of tumors. A clustering of the proteomic profiles of the samples visually confirmed this, as the tumor profiles clearly display a greater heterogeneity than the controls (F test p - value: 3e - 6, Fig. 1B, S1D). The minimum correlation between pairs of biological replicates ranged from 0.75 to 0.95 (Fig. S1D). A differential expression analysis was performed between the control and tumor samples using linear models in order to estimate the significance of the changes in protein abundances (tumor / control, Fig. 1C and Materials and Methods). Out of 4.415 proteins, 1.886 had a significant fold change and 1.018 were associated with a False Discovery Rate (FDR) < 5%. The log2 fold change appears to be symmetrically distributed around zero. 1.263 (28.6%) out of the 1.996 tested protein uniprot identifiers were associated with the GO term metabolic process; 702 of these had a significant fold change (FDR < 0.05, Fig. 1D). Thus, since the ratios tested/significant (FDR 0.05) proteins and tested/significant metabolic associated proteins are similar (FDR 0.53 and 0.55), global protein abundance changes demonstrates that the majority of differentially expressed proteins is associated with metabolism (68.9% of all FDR - significant proteins). However, metabolism might be targeted preferentially by directional regulations. Hence, gene set analysis was performed using the PIANO package (Varemo et al., 2013), in order to find significant directional alterations of metabolism - related pathways by incorporating fold - change directionality in the statistical enrichment analysis. Many central metabolic pathways were found to be significantly down - regulated (FDR ≤ 0.05), such as oxidative phosphorylation, citrate cycle and fatty acid metabolism (Fig. 1E, S2). This observation suggested a robust reprogramming of cellular metabolism in the tumor samples, especially towards a down - regulation of the abundance of proteins involved in energy metabolism.

**Figure 1.**
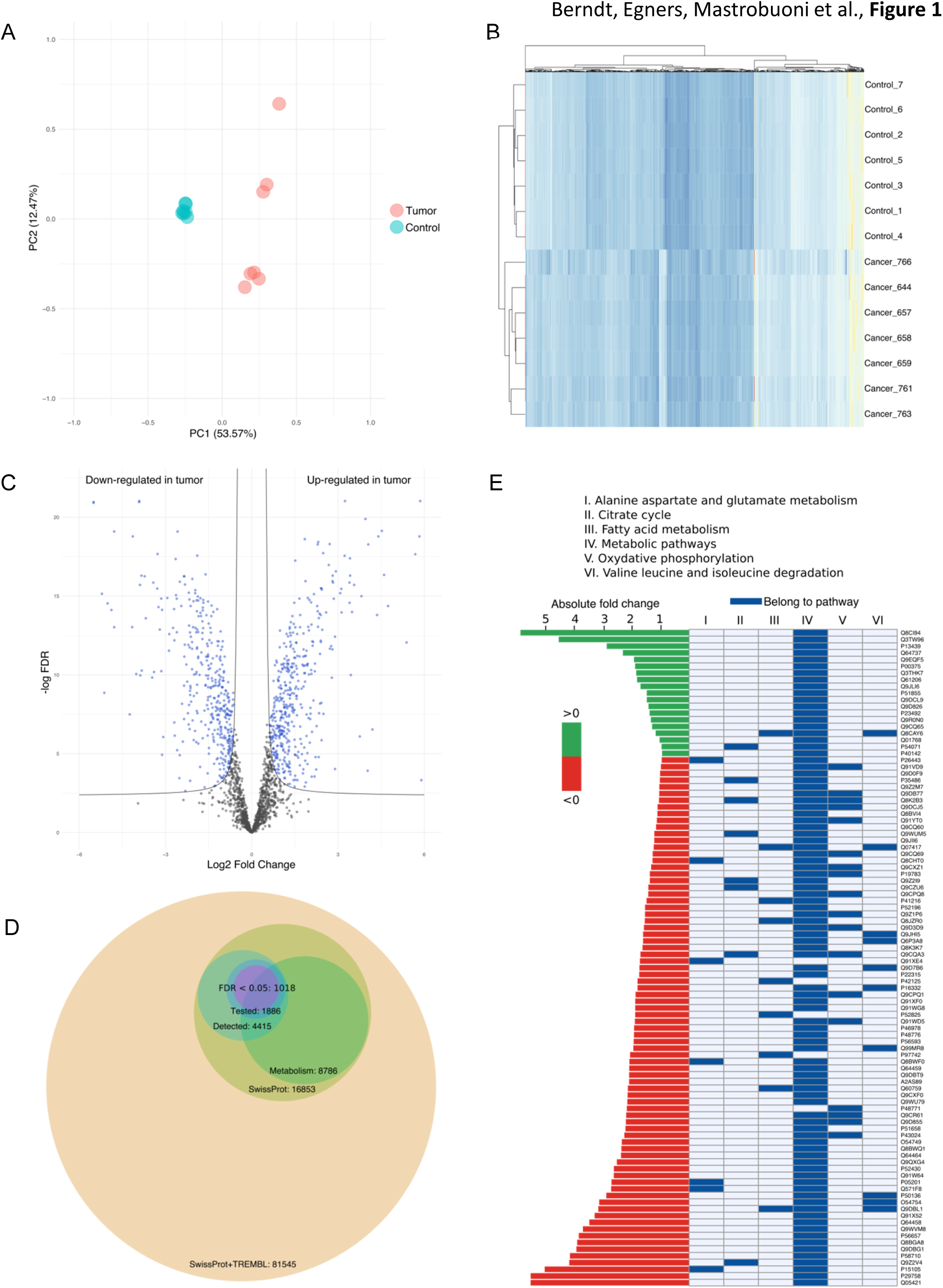
Detected metabolic enzymes in normal and HCC mouse liver and data quality control. (A) Principal Component Analysis (first two components, 53.57% + 12.4% of variance). Control (blue) and tumor (red) samples are well separated on the first component. (B) Clustering of the complete cases of proteomic samples. Control and tumor samples cluster together, respectively. (C) Volcano plot showing the log2 fold changes of proteins (tumor/control) with respect to the - log of FDR. Left side correspond to proteins that are down - regulated in tumor, while right side correspond to proteins that are up - regulated in tumor. (D) Bubble plot showing the relations between the different protein sets considered in the study. Out of the 16.853 reviewed proteins present in the SwissProt database (of which 8.786 are associated with metabolism), 4.415 were identified by mass spectrometry. Significance of the fold changes between tumor and control could be estimated for 1.886 proteins, of which 1.018 passed the threshold of 5% FDR. (E) Histomap showing the highly significant fold changes of 145 proteins (FDR ≤ 0.0001) associated with 6 significantly down - regulated metabolic pathways (FDR ≤ 0.05, protein sampling).

In particular, we found that the majority of glycolytic enzymes such as PFK - L (the liver - specific isoform of 6 - phosphofructokinase) and GAPDH (glyceraldehyde - 3 - phosphate dehydrogenase) are significantly upregulated in HCC tissues (Fig. S3). Furthermore, the fructose - bisphosphate aldolase isoforms A and C, which - unlike isoform B - preferentially contribute to glycolytic rather than gluconeogenic metabolite turnover, show a more than two - fold higher expression. In contrast, enzymes of other important metabolic pathways are downregulated such as pyruvate carboxylase, citrate synthase, succinate dehydrogenase, carnitine O - palmitoyltransferase 2, glutaminase (liver isoform), glutamine synthetase and ornithine carbamoyltransferase (Fig. S3).

### Prediction of tumor - specific metabolic capacity via mathematical modeling

Relative changes of protein abundances were mapped onto the maximal capacities of the respective enzymes to generate a kinetic model of the central metabolism of murine HCC (Fig. 2). To assess the functional consequences of alterations in metabolic enzyme expression, we applied the model to a typical 24h physiological plasma concentration profile of exchangeable model metabolites and the hormones insulin and glucagon. We used the plasma profile as input for the model and computed the diurnal variations in the concentrations of all model metabolites and fluxes for normal liver and murine HCC (Fig. 3). Compared with normal hepatocytes, the simulated metabolic response of murine HCC revealed a number of significant alterations (Fig. 3). The activity of glycolysis is strongly elevated while gluconeogenesis is almost completely suppressed. In line with these alterations, HCC is predicted to operate continuously as a strong lactate producer, while normal hepatocytes take up lactate. Fatty acid uptake, ß - oxidation of fatty acids, fatty acid and cholesterol synthesis are strongly diminished. Oxygen consumption is lower in HCC compared to normal liver. In addition, ammonia detoxification and urea synthesis in HCC are also rigorously reduced (Fig. 3).

**Figure 2.**
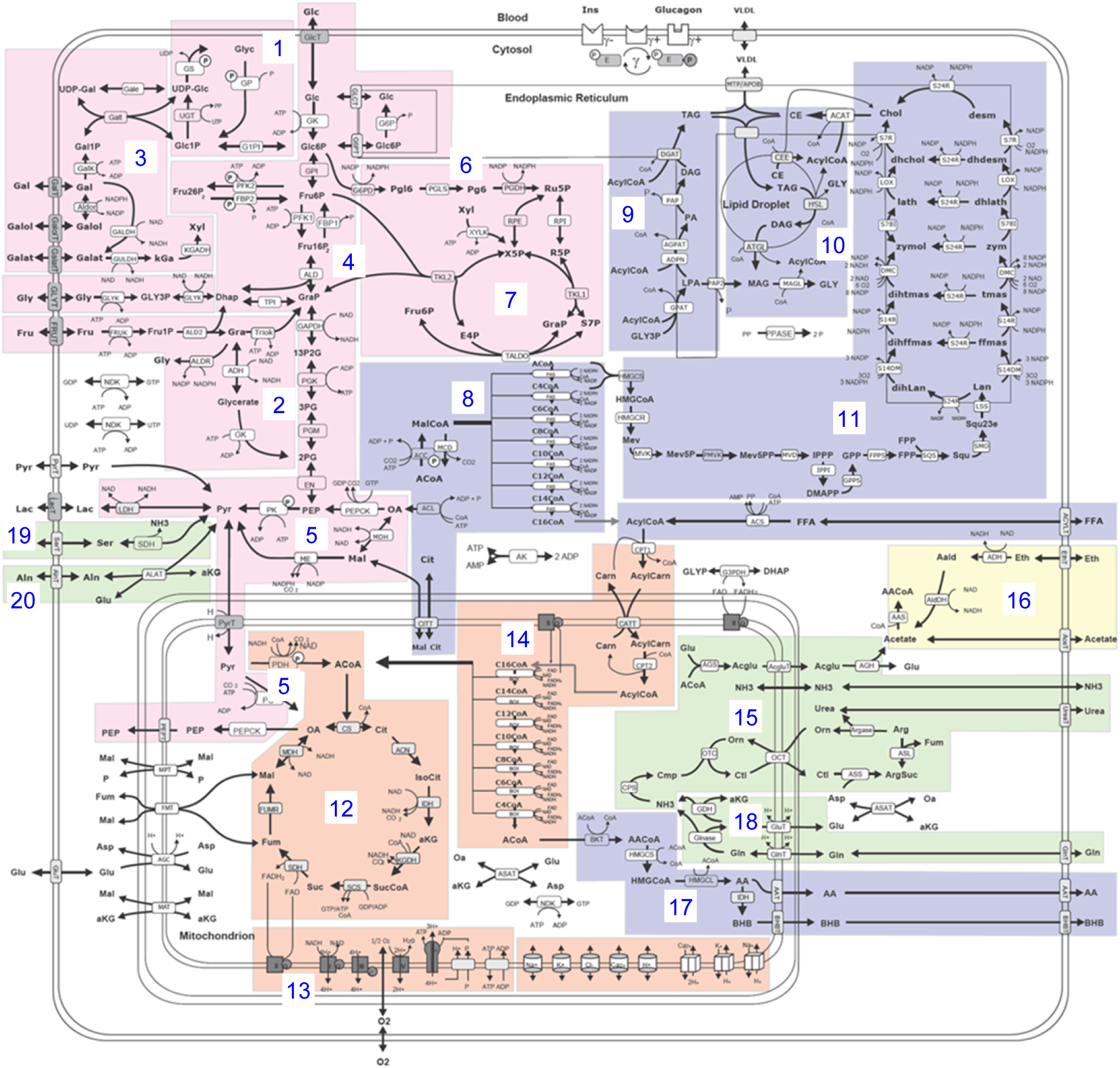
Central carbon metabolism pathways covered by the kinetic model. Glycogen, fructose, galactose metabolism (1, 2, 3), glycolysis (4), gluconeogenesis (5), oxidative and non - oxidative pentose phosphate pathway (6, 7), fatty acid and triglyceride synthesis (8, 9), synthesis and degradation of lipid droplets (10), cholesterol synthesis (11), TCA cycle (12), respiratory chain and oxidative phosphorylation (13), β- oxidation (14), urea cycle (15), ethanol metabolism (16), ketone body synthesis (17), glutaminolysis and glutamine synthesis (18), serine and alanine utilization (19, 20).

**Figure 3.**
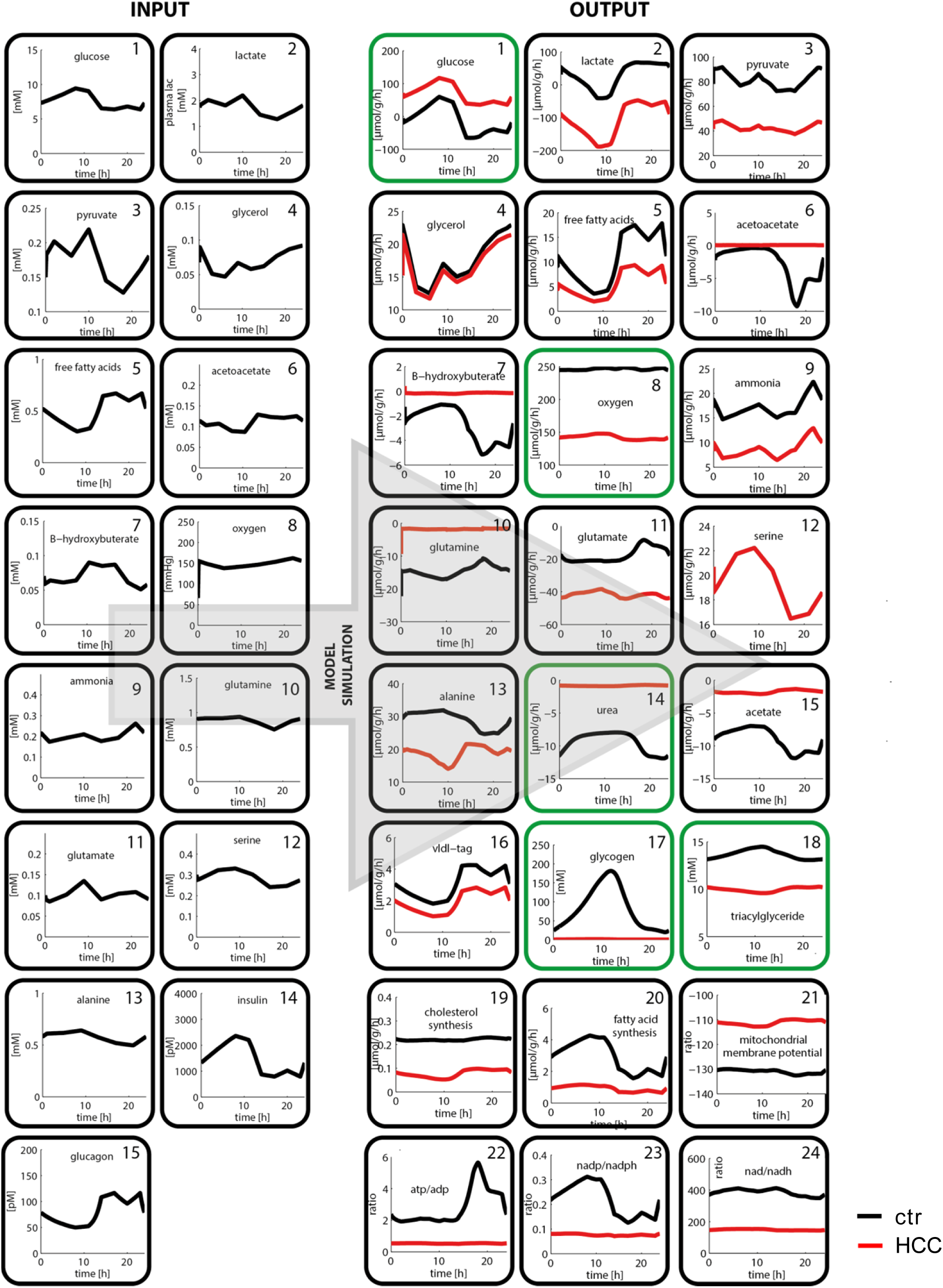
Input and output parameters of the metabolic model. Experimentally validated model predictions are highlighted in green.

### Experimental validation of model predictions

Next, we sought to perform a functional validation of selected model predictions. To validate the predicted changes of glycolysis (Fig. 4A) and mitochondrial function (Fig. 4E), extracellular flux analyses of normal primary hepatocytes (isolated from healthy C57Bl/6J mice) and isolated HCC cells (from ASV - B mice) were performed. As can be seen in figure 4B, the extracellular acidification rate (ECAR) as a quantitative read - out of glycolytic activity is significantly elevated in HCC cells (t - test, p<0.0001). Moreover, in contrast to hepatocytes, HCC cells are capable of further increasing the glycolytic rate after inhibition of mitochondrial respiratory chain complexes. In line with these results, we found that HCC cells isolated from ASV - B mice grow significantly slower in medium without glucose compared to standard medium (25 mM glucose, Fig. 4C; t - test, p<0.05). Pulsed stable isotope - resolved metabolomics (pSIRM) revealed higher label incorporation into lactate after intraperitoneal administration of ^13^C - glucose by ASV - B tumors compared to normal liver (Fig. 4D). This argues for elevated glycolytic activity of murine HCCs, well in line with the above outlined model prediction (Fig. 4A). To test the mathematically predicted changes in oxygen uptake (Fig. 4E), the oxygen consumption rate (OCR) was determined and found to be decreased in HCC cells (Fig. 4F; t - test, p<0.0001). In addition, calculation of the OCR/ECAR - ratio showed that primary hepatocytes prefer oxidative phosphorylation over glycolytic energy production to a significantly greater extent than their HCC counterparts (Fig. S4; t - test, p<0.0001). To further analyze this, we quantified the cellular mitochondrial content with electron microscopy. As can be seen in figure 4G and supplementary figure 4B, these analyses indicated that the number of mitochondria is indeed significantly different between normal liver and murine HCC. Taken together, these functional assays display reduced mitochondrial activity in murine HCC and hence nicely confirm the model prediction shown in Figure 4E. As outlined above, model calculations reveal a diminished capacity of HCC tumor tissue to synthesize urea in order to detoxify ammonia (Fig. 5A). We validated this prediction by measuring the urea concentration in the cell culture supernatant and indeed found significantly less urea in the supernatant of ASV - B cells (Fig. 5B; t - test, p<0.0001). Comparing the amount of urea produced by precision cut liver slices (PCLS) from normal and HCC liver supports this result (Fig. 5C; t - test, p<0.0001). Additionally, we challenged the model prediction of impaired triacylglyceride production capacity in HCC cells (Fig. 5D) by analyzing the intracellular amount of triacylglycerides of cells without and after supplementation of the culture medium with oleic acid. Triacylglycerides were readily detectable in primary hepatocytes and their amount was increased by providing oleic acid. In contrast, no triacylglycerides were detectable in ASV - B cells regardless of the presence or absence of oleic acid (Fig. 5E). One key function of the liver is the intracellular storage of glycogen which was predicted to be completely abolished in HCC tumors (Fig. 6A). By performing Periodic acid – Schiff (PAS) staining for glycogen detection on sections from normal and ASV - B liver we in fact saw diminished staining in murine HCC tumors (Fig. 6B).

**Figure 4.**
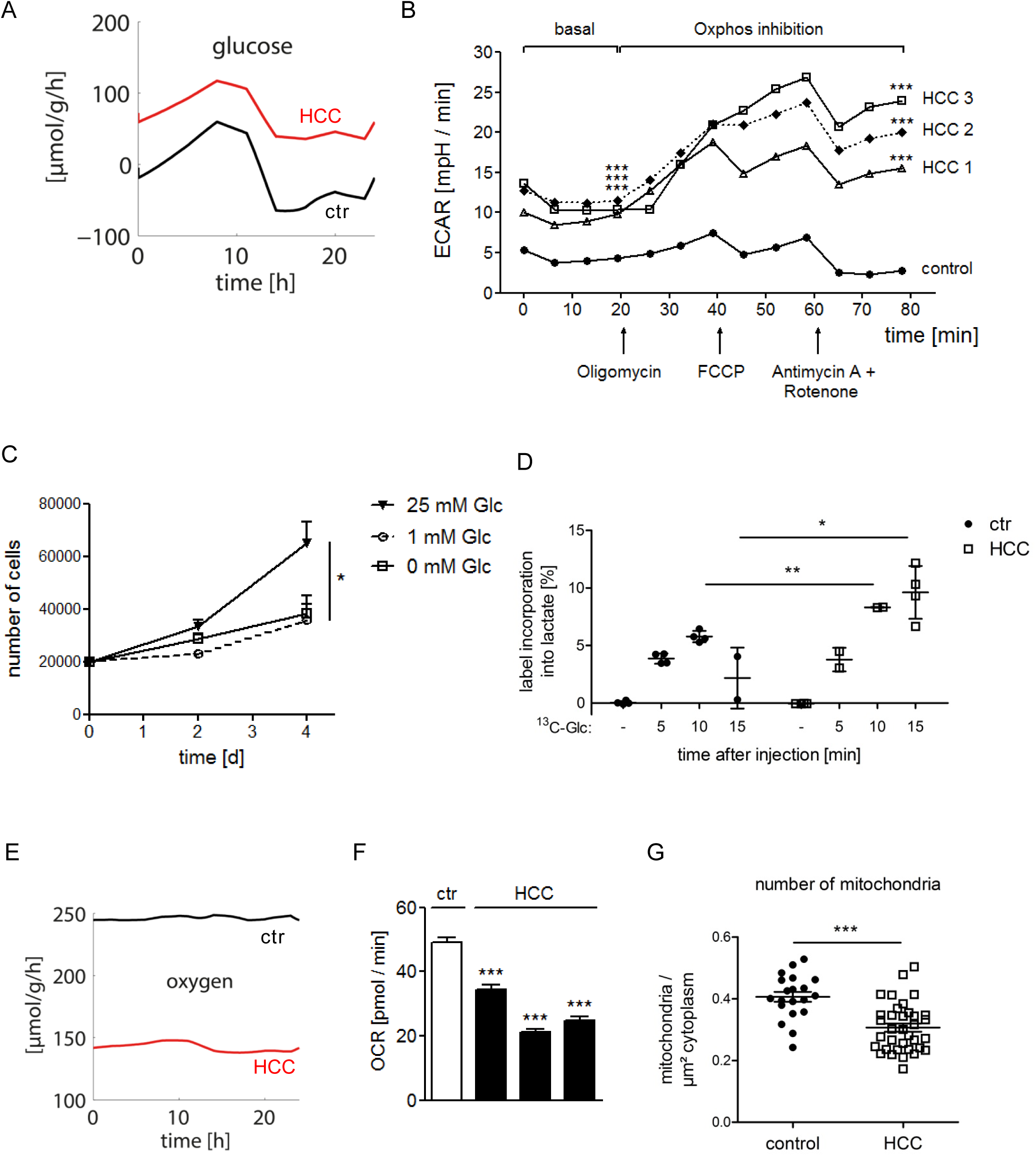
Experimental verification of predicted elevated glycolytic activity and reduced oxygen consumption in murine HCC. (A) Simulating the glucose exchange flux shows complete impairment of glucose secretion and higher glucose uptake by tumors compared to control liver. (B) Basal and post - respiratory chain complex inhibition extracellular acidification rates (ECAR) of isolated HCC cells and primary hepatocytes were measured. (n = 10). (C) Varying media glucose concentrations affect the proliferation of isolated HCC cells. (n = 3). (D) *In vivo* pSIRM experiments reveal higher ^13^C incorporation into lactate in tumors after i.p. injection of ^13^C - glucose. (E) Kinetic model calculations and metabolic flux analysis on isolated cells (F) show a lowered oxygen consumption rate of HCC cells. (G) The number of mitochondria per cell was quantified by electron microscopy. (*, p < 0.05, **, p < 0.005; ***, p < 0.0001).

**Figure 5.**
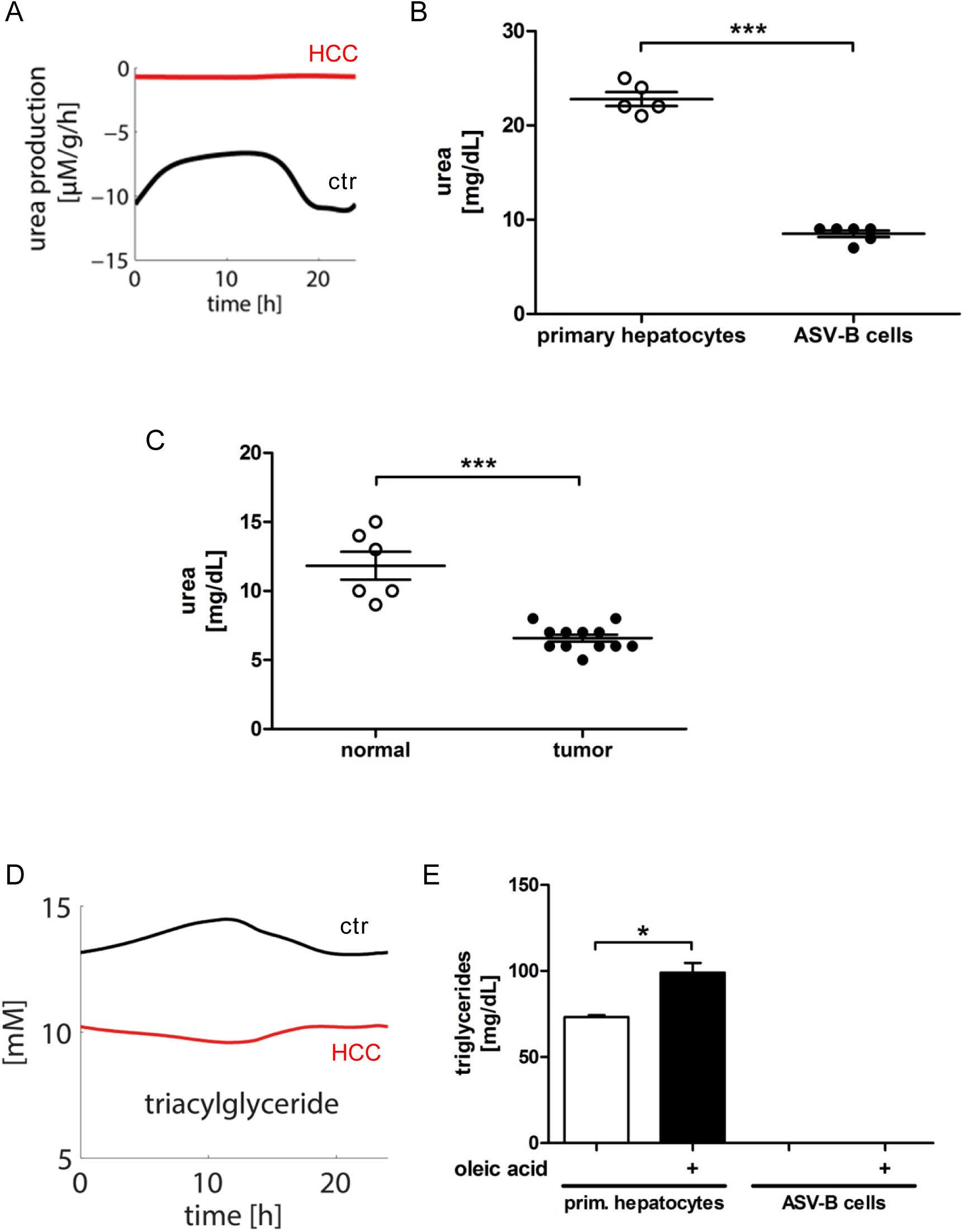
Experimental validation of urea production and intracellular triacylglyceride. (A) Model calculations predict the inability of HCC tumors to produce urea. (B) Determining the urea concentration in the supernatant of isolated hepatocytes and ASV - B cells and (C) PCLS. (D) Model simulations show a reduced capacity of HCC tumors to synthesize triacylglyceride. (F) Measurement of intracellular triacylglyceride in primary hepatocytes and isolated HCC cells without and after addition of oleic acid into the culture medium. (*, p < 0.05; ***, p < 0.0001).

**Figure 6.**
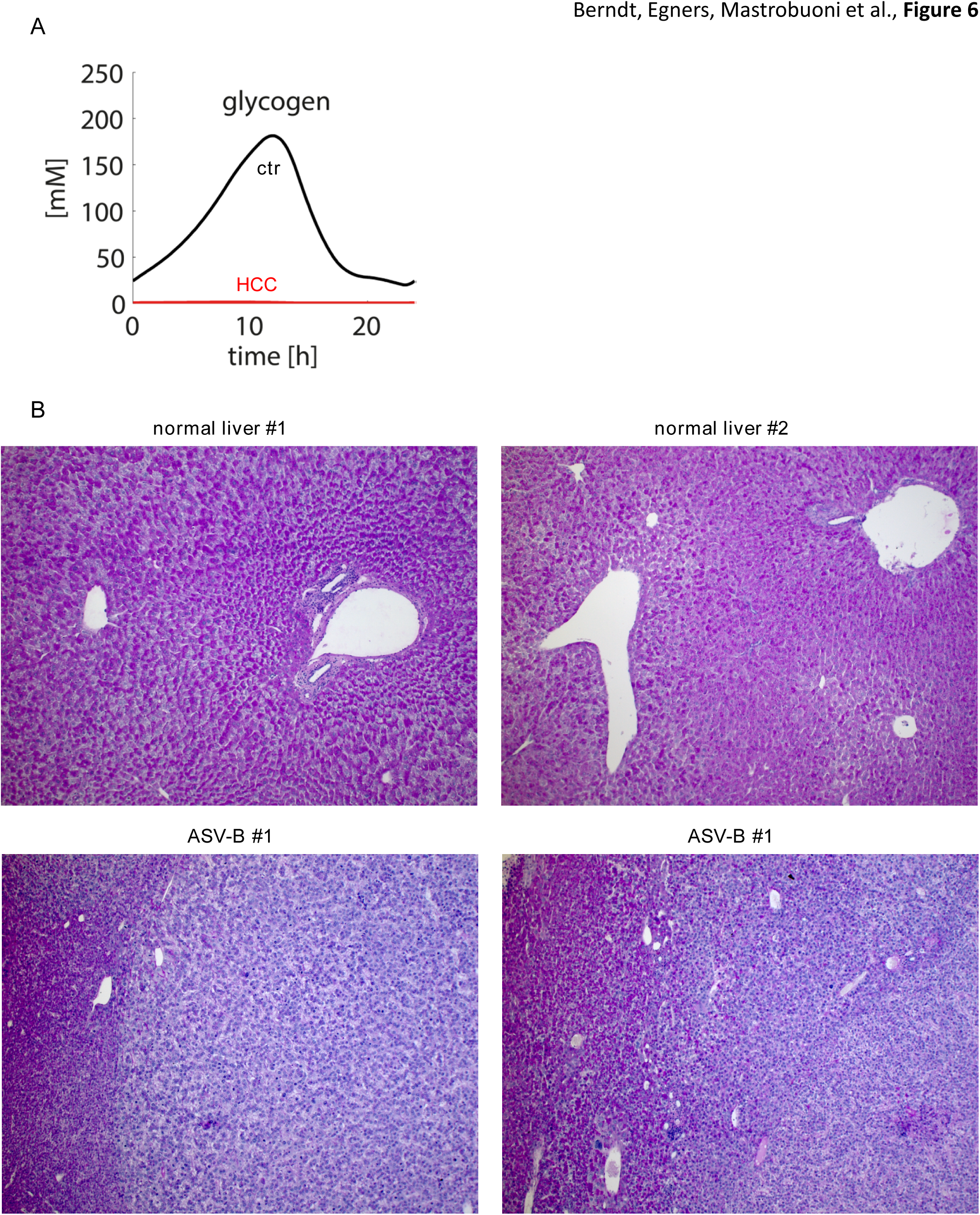
ASV - B HCC tumor cells produce and store less intracellular glycogen. (A) According to the model prediction, HCC tumors do not store any glycogen. (B) PAS staining for intracellular glycogen of normal and ASV - B liver sections.

### Model - based predictions and functional validation of tumor - specific metabolic vulnerabilities

We hypothesized that the reduced capacity of oxidative phosphorylation in HCC can be exploited for cancer therapy by serving as a metabolic target to selectively impair HCC metabolism while leaving healthy liver intact. This was further strengthened by modeling the oxygen consumption as a function of mitochondrial complex I activity, demonstrating a greater sensitivity of tumor compared to control liver (Fig. 7A). The antidiabetic drug metformin has been shown to inhibit neoplastic growth by multiple mechanisms (Coyle et al., 2016), one of them being complex I inhibition (Wheaton et al., 2014). In addidtion, it was shown that metformin acts as non - competitive inhibitor of mitochondrial glycerophosphate dehydrogenase (Mgpdh), explaining its antidiabetic properties (Madiraju et al., 2014). Using the reported inhibition constants of metformin of 0.5 mM for complex I (Wheaton et al., 2014) and 0.055 mM for Mgpdh (Madiraju et al., 2014), we simulated the effect of metformin on HCC and healthy liver. We put the external conditions to their mean value over one day and varied the metformin concentration from 0 to 1 mM. Figure 7B depicts the mitochondrial membrane potential of healthy liver and murine HCC as a function of the metformin concentration. As mitochondria induce apoptosis in response to energy depletion (once the mitochondrial membrane potential falls below ∽ - 80 mV), the simulations predict damage to the liver tumors already at 0.27 mM metformin, while healthy hepatocytes remain viable up to metformin concentrations of 0.7 mM. We functionally validated these results by treating PCLS with 0.5 mM metformin. According to our model’s predictions, exposure to metformin did not affect healthy liver tissue (Fig. 7C), but resulted in a significant increase of HCC cells undergoing cell death (Fig. 7D; t - test, p<0.05).

**Figure 7.**
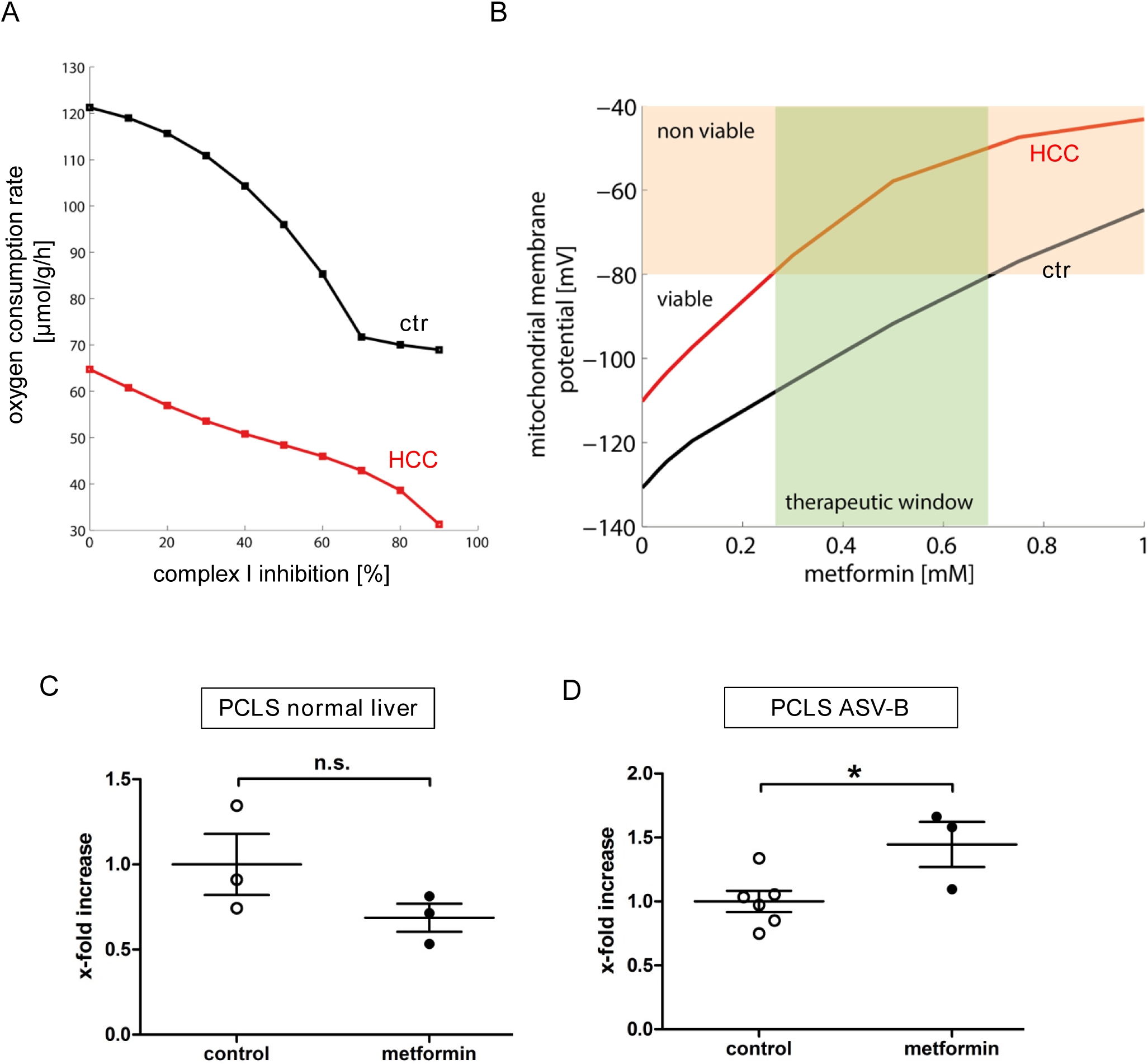
Mathematical sensitivity analysis identifies complex I inhibition as an effective anti - proliferative treatment for murine HCC. (A) Oxygen consumption rate of control and HCC tumor liver under the condition of complex I inhibition and (B) calculated mitochondrial membrane potential. (C, D) Analysis of LDH release into the supernatant after metformin treatment of PCLS from normal and ASV - B mice (*, p < 0.05).

## Discussion

The fundamental metabolic reprogramming processes that tumor cells undergo to support growth and survival have received widespread attention in recent years and are now considered as an emerging hallmark of cancer (Hanahan and Weinberg, 2011; Pavlova and Thompson, 2016). However, exploitation of metabolic vulnerabilities to identify effective and specific anti - cancer agents remains challenging. The advancement of analytical technologies like shotgun proteomics opened the way for global snapshots of the molecular makeup of healthy tissues and tumors (Uhlén et al., 2015; Yu et al., 2016). The increased sensitivity of these technologies together with improved data reproducibility enable for the first time to map the biochemical network in its totality. However, due to (i) enormous plasticity and dynamics, (ii) multi - level regulatory mechanisms and (iii) a highly complex network of reactions, cellular metabolic processes are difficult to study. Mathematical models are useful tools to unravel this complexity, and various tools have been established already to simulate metabolism and to predict metabolic activities of cells (Berndt and Holzhutter, 2016). Naturally, mathematical models are always simplifications of multi - level biological phenomena; however, hitherto published metabolic models of liver metabolism specifically lack important regulatory aspects, e.g. hormonal influences and allosteric parameters. In addition, they are very often not based on data that have been established experimentally but on information solely extracted from published literature.

Here, we used a comprehensive kinetic model of the central carbon and lipid metabolism of hepatocytes that incorporates not only the metabolic reaction network but also enzyme regulation by allosteric effectors and by reversible phosphorylation due to changing insulin and glucagon signaling (Berndt *et al.*, under review). The influence of fluctuating nutrient (like glucose and glutamine) and oxygen concentrations within a physiological range are also taken into account. As demonstrated by us earlier, these parameters are at least equally important for modelling the metabolic performance as the changes in enzyme abundance (Bulik et al., 2016). Applying the model to the central metabolism of HCC, we took advantage of the fact that HCC tumors originate from hepatocytes (Mu et al., 2015), i.e. metabolic enzymes in normal and malignant cells only differ in their expression level. This enabled us to re - parameterize the hepatocyte model by scaling the enzyme activities between HCC and normal hepatocytes according to the observed changes of protein abundances that we assessed by comprehensive mass spectrometry. Our study proofs that this approach is able to transform mass spectrometry protein data into biologically and clinically meaningful metabolic predictions about the HCC tissue turn - over activity of glucose, lactate, pyruvate, glycogen, fatty acids, ammonia, urea, amino acids and oxygen, to mention only some. Comparison of simulated normal and HCC liver metabolic performance reveals fundamental differences. We validated several of the calculated model output parameters successfully using different experimental approaches. Glycolytic and respiratory activity were determined by measuring the extracellular acidification and oxygen consumption rates, confirming that HCC cells show increased glycolytic and reduced mitochondrial activity. These findings are well in line with earlier reports using transcriptomics, metabolomics or enzyme activity measurements on different murine HCC models (Bard - Chapeau et al., 2014; Dolezal et al., 2017; Fan et al., 2017). Increased glycolytic activity is consistently found in independent analyses of human HCC tissue with different omics approaches and non - invasive imaging (NMR spectroscopy), pointing towards the Warburg effect as a metabolic hallmark of human liver cancer (Beyoglu et al., 2013; Budhu et al., 2013; Yang et al., 2007).

The kinetic model predicted HCC - specific alterations of urea and triacylglyceride synthesis as well as glycogen storage, all of which represent key functions of normal liver. The functionality of the urea cycle in HCC tissue has been under debate for quite some time. It had been established rather early on that HepG2 cells, one of the most widely used human HCC cell lines, harbor a defective urea cycle and it was later shown that this is due to ornithine transcarbamylase and arginase I deficiency (Mavri - Damelin et al., 2007). On the other hand, arginase I and carbamoyl phosphate synthetase (CPS) were found overexpressed in human HCC and their detection via immunohistochemistry was demonstrated to improve the histopathological diagnosis of HCC (Butler et al., 2008; Yan et al., 2010). Our approach now reveals - for the first time - reduced urea cycle activity in a murine HCC model, nicely confirming that systematic integration of protein expression data is a prerequisite to comprehend metabolic pathway activity (Berndt and Holzhutter, 2016). The analysis of HCC - specific changes of lipid metabolism has received a lot of attention recently (over 20 studies using human samples and different omics approaches were published in the last 10 years (Beyoglu and Idle, 2013)). The reported results are very heterogeneous, precluding the identification of a HCC - specific lipid metabolism pattern. If anything, activation of fatty acid catabolism (most importantly β- oxidation) could be considered a hallmark of HCC - specific lipid metabolism as it was reported by the majority of publications (Beyoglu and Idle, 2013). Of note, the activity of anabolic lipid metabolism pathways in HCC has received significantly less attention. Our approach of combining quantitative proteomics with mathematical modeling predicted reduced activity of several anabolic lipid pathways in HCC, e.g. synthesis of triacylglycerides, cholesterol and fatty acids as well as VLDL secretion (Figure 3). We were able to functionally validate the calculated reduction of triacylglyceride synthesis, underscoring the eligibility of the comprehensive kinetic model to forecast alterations of lipid metabolism in HCC. While our results predicted and validated reduced glycogen storage in murine HCC, none of the above mentioned omics analyses of human HCC reported reduced glycogen concentrations (Beyoglu and Idle, 2013). This is especially intriguing as it has been known for quite some time that human HCCs differ with respect to the extent of the Periodic acid - Schiff (PAS) reaction, the routine histochemical detection method for glycogen (and other complex carbohydrates) (Kitamura et al., 1993). These differences are clinically relevant as the survival of patients with PAS - negative HCCs was significantly shorter than that of PAS - positive ones (Kitamura et al., 1993). Whether this observation is functionally linked to altered glycogen metabolism has not been addressed thus far. One could hypothesize that glycogen storage and the Warburg effect are inversely correlated in HCC. With rising malignancy, all available glucose is needed for the Warburg effect (to enable neoplastic proliferation (Pavlova and Thompson, 2016)), resulting in a functional loss of glycogen storage. This, in turn, would translate into reduced ability of the tumor cells to secrete glucose and hence to participate in glucose homeostasis, another key function of the normal liver. As can be seen in figure 3, our kinetic model predicts a complete loss of glucose secretion by murine HCC tissue, supporting the above outlined hypothesis. In summary, the results of all validation experiments show striking consistency with the calculated model simulations indicating that our kinetic model is indeed a powerful tool to reproduce HCC metabolism in a reliable manner.

The feasibility of the translational application of our kinetic model to estimate and evaluate the performance of therapeutic agents with prediction of possible adverse effects is demonstrated by calculating the outcome of metformin treatment on HCC viability. The anti - diabetic drug metformin received a lot of attention in recent years after it was reported to reduce cancer risk and mortality in diabetic patients (Evans et al., 2005). Metformin was subsequently shown to exert antitumor effects against established human HCC cell lines and in HCC xenografts in nude mice (Miyoshi et al., 2014). Our results confirm the cell line data reported by Miyoshi *et al.* and furthermore show that primary hepatocytes are not affected by metformin at the doses found to inhibit HCC cells. These results demonstrate the versatility of our kinetic model as – in addition to depicting metabolic activities - it is able to predict metabolic vulnerabilities that can potentially be exploited for cancer therapy.

In the future, intensive effort has to be invested into the advancement of comprehensive kinetic model systems in order to further approximate to the essential processes in pathologic tissues and to increase the quantity of accomplishable model calculations. Since the metabolism of cells and tissues is embedded in and influenced by a large network of additional interconnected reactions which can be studied by comprehensive proteomics analyses, the expansion of the included model framework by parameters other than metabolic ones is certainly required in order to generate a global scale mathematical model characterizing a respective tissue. In this way the prospective utilization of kinetic models in personalized medical care in which every patient receives a treatment whose effects and unwanted side effects have been simulated and consequently evaluated beforehand will transform into a conceivable and concrete vision.

## Author contributions

NB, AE, GM, OV, AF, RvG, SOD, MP, ME, HGH, SK and TC performed and/or coordinated experimental work. NB, AE, GM, AF, AD, MP, CB, RvG, ME, JSR and SK performed data analysis. AE, AF, ME and TC performed animal experiments. NB, AE, GM, AD, JSR, HGH, SK and TC prepared the initial manuscript and figures. HGH, SK and TC provided project leadership. All authors contributed to the final manuscript.

## Acknowledgements

This work was funded by a grant from the Bundesministerium für Bildung und Forschung (BMBF) to Thorsten Cramer, Stefan Kempa and Hermann - Georg Holzhütter (HepatomaSys in the framework of “e:Bio - Innovationswettbewerb Systembiologie”). Research in the Cramer laboratory was further supported by grants from the Deutsche Forschungsgemeinschaft (DFG) and the Deutsche Krebshilfe. Stefan Kempa received funding by the Senate of Berlin, the Berlin Institute of Medical Systems Biology (BIMSB) and the Berlin Institute of Health (BIH). The Q3 European Union Horizon 2020 grant SyMBioSys (MSCA - ITN - 2015 - ETN #675585) provided financial support for Aurélien Dugourd.

## Materials and Methods

### Transgenic HCC Model and tissue preparation

The murine HCC model (termed ASV - B) was established and initially characterized by Dubois and co - workers (Dubois et al., 1991). Briefly, male ASV - B mice express the early region of the SV40 large T (SV40lT) oncogene under control of the mouse antithrombin III promoter. ASV - B mice show time - dependent liver tumor development with first evidence of dysplasia at 8 weeks, adenomas at 12 weeks and hepatocellular carcinoma (HCC) at 16 weeks of age. All mice were maintained under standard conditions at the animal facilities in Berlin (Charité) and Aachen (Institut für Versuchstierkunde, University Hospital Aachen). Animal procedures were performed in accordance to approved protocols (Landesamt für Gesundheit und Soziales Berlin (0024/12) and Landesamt für Natur, Umwelt und Verbraucherschutz Düsseldorf (84 - 02.04.2015.A344, AZ84 - 02.04.2016.A018 and 84 - 02.04.2015.A216)) and followed recommendations for proper care and use of laboratory animals. For tissue preparation, 16 weeks old ASV - B or tumor - free male control mice (C57Bl/6J, Harlan Laboratories) were sacrificed by cervical dislocation and liver tissue samples were snap - frozen in liquid nitrogen for further analysis.

### Proteome analysis

Murine liver samples were immediately frozen in liquid nitrogen and resuspended in a urea buffer (8 M urea, 100 mM TrisHCl, pH 8.25) containing 100 μl of zirconium beads for protein extraction. Samples were homogenized on a Precellys 24 device (Bertin Technologies) for two cycles, 10 seconds at 6,000 rpm. After centrifugation to remove beads and tissue debris, protein concentration was measured by Bradford colorimetric assay and 100 μg were taken for protein digestion. Leftover samples were frozen at - 80°C. The disulfide bridges of proteins were reduced in DTT 2 mM for 30 minutes at 25°C and successively free cysteines alkylated in iodoacetamide 11 mM for 20 minutes at room temperature in the dark. LysC digestion was then performed by adding 5 μg of LysC (Wako Chemicals) to the sample and incubating it for 18 hours under gentle shaking at 30°C. After LysC digestion, the samples were diluted 3 times with 50 mM ammonium bicarbonate solution, 7 μl of immobilized trypsin (Applied Biosystems) were added and samples were incubated 4 hours under rotation at 30°C. 18 μg of the resulting peptide mixtures were desalted on STAGE Tips (Rappsilber et al., 2003) and the eluates dried and reconstituted to 20 μl of 0.5 % acetic acid in water.

### LC - MS/MS analysis

5 μl were injected in duplicate on a UPLC system (Eksigent Technologies, USA), using a 240 minutes gradient ranging from 5% to 45% of solvent B (80% acetonitrile, 0.1 % formic acid; solvent A = 5 % acetonitrile, 0.1 % formic acid). For the chromatographic separation 30 cm long capillary (75 μm inner diameter) was packed with 1.9 μm C18 beads (Reprosil - AQ, Dr. Maisch HPLC, Germany). On one end of the capillary nanospray tip was generated using a laser puller, allowing fretless packing. The nanospray source was operated with a spay voltage of 2.1 kV and an ion transfer tube temperature of 260°C. Data were acquired in data dependent mode, with one survey MS scan in the Orbitrap mass analyzer (60,000 resolution at 400 m / z) followed by up to 20 MS / MS scans in the ion trap on the most intense ions. Once selected for fragmentation, ions were excluded from further selection for 30 seconds, in order to increase new sequencing events.

### *In vivo* pulsed stable isotope resolved metabolomics

Frozen liver tissue was grounded using mortar and pestle and the powdered tissue was directly resolved in pre - cooled extraction buffer (- 20°C, chloroform/methanol/water (2:5:1 vol / vol / vol). 50 mg tissue were resolved in 1 ml extraction buffer including the internal standard cinnamic acid (Sigma - Aldrich) 2 μg/ml. Samples were shaken for 1 hour at 4°C, subsequently 500 μl water were added, samples shaken for 30 minutes at 4°C, centrifuged for 10 minutes at 10,000g and polar and lipid phase collected and dried overnight in a speed vac.

### Metabolomics measurements and data analysis

Dried polar extracts were resolved in 20 μl pyridine including 40 mg/ml Methoxamine hydrochloride (Sigma - Aldrich) and shaken for 1 hour at 30°C. Subsequently, 80 μl Methyl - N - (trimethylsilyl) trifluoroacetamide (MSTFA, Fluka) was added and samples were shaken for 90 min at 37°C. Samples were centrifuged (10 min at 10,000g) and transferred into glass vials (Chromacol). 1 μl of the samples was injected into an Agilent gas chromatograph 6890 equipped with a temperature controlled injection system (ALEX, Gerstel). Data were acquired using a GC - ToF - MS (Pegasus III, Leco), processed using the vendor software Chromatof (Leco) and analyzed using the MetMax software (Kempa et al. 2009 (PMID: 19206143)). Identification of metabolites was performed using pure chemicals (Sigma - Aldrich), quantification of metabolites and calculation of isotope incorporation were performed as described (Pietzke et al., 2014).

### Metabolic model

We have recently developed a comprehensive kinetic model of the central metabolism of hepatocytes (Berndt *et al.*, manuscript under review). The model was used to simulate the impact of nutrient supply (including oxygen), hormonal stimuli and protein abundance of metabolic enzymes on the functional output of the liver. The model comprises the central hepatic metabolic pathways of glycolysis, gluconeogenesis, glycogen synthesis, glycogenolysis, fructose metabolism, galactose metabolism, the creatine - phosphate/ATP shuttle system, the pentose phosphate cycle composed of the oxidative and non - oxidative branch, the citric acid cycle, the malate aspartate redox shuttle, the glycerol - 3 - phosphate redox shuttle, the mitochondrial respiratory chain, the beta - oxidation of fatty acids, fatty acid synthesis, ketone body synthesis, cholesterol synthesis, triglyceride synthesis and degradation, the synthesis and hydrolysis of triglycerides, the synthesis and export of the very - low density lipoprotein (VLDL), the urea cycle, the metabolism of the amino acids serine, alanine, glutamate, glutamine, aspartate and ethanol metabolism. The model contains the key electrophysiological process of the inner mitochondrial membrane including the mitochondrial membrane potential, mitochondrial ion homeostasis and the generation and utilization of the proton motive force. The modeled reactions and transport processes are depicted in figure 2. The metabolic model is coupled to a phenomenological model of hormonal signaling by glucagon and insulin affecting the short - term regulation of metabolic enzymes by reversible phosphorylation (see below). The model describes the uptake, metabolization and generation of glucose, fructose, galactose, pyruvate, lactate, glycerol, ammonia, serine, alanine, glutamate, glutamine, fatty acids, ethanol, acetate, urea, acetoacetate, β - hydroxybutyrate, oxygen and VLDL particles.

### Short - Term Regulation of Liver Metabolism by Hormones

The metabolism of the liver is strongly controlled by hormones, in particular insulin and glucagon (Ohno and Maier, 1994). Glycolysis and gluconeogenesis as well as fatty acid synthesis and β- oxidation are inversely regulated by glucagon and insulin signaling via phosphorylation and de - phosphorylation of key regulatory enzymes. In the model, the plasma concentration of insulin and glucagon is directly translated into the phosphorylation state of interconvertible enzymes by a phenomenological sigmoid function (γ - function) also used in (Bulik et al., 2016). Moreover, we used phenomenological transfer functions (Fig. 2) to compute the plasma concentrations of insulin and glucagon and of non - esterified fatty acids (NEFA) directly from the plasma level of glucose. This setting rests on the assumption that the release of insulin and glucagon from pancreatic islet cells is mainly controlled by the plasma glucose level and that high concentrations of glucagon and epinephrine stimulate the hormone - sensitive lipase (HSL) in adipose tissues, thus creating an inverse relationship between the plasma level of glucose and NEFA.

### Isolation and culture of primary murine hepatocytes and establishment of ASV - B cell lines

Primary hepatocytes from C57Bl/6J were isolated as described earlier (Hesse et al., 2012) and maintained in Dulbecco’s modified Eagle’s medium (DMEM) containing 10% fetal bovine serum and 1% penicillin - streptomycin (Life Technologies) in collagen - coated flasks. All experiments were conducted within five days after primary cell isolation. HCC cells were isolated from 16 weeks old ASV - B mice and cultivated in the medium described above. After an initial adaptation period of one to two months they started to proliferate and grow stably under cell culture conditions.

### Cellular metabolic rate measurements

Primary hepatocytes and ASV - B cells were seeded 24 hours ahead of analysis (7,000 cells per well in collagen - coated XF96 cell culture plates, Seahorse Bioscience) in standard medium. One hour before experimental start the cells were supplied with XF Base medium (Seahorse Bioscience) supplemented with 2 mM glutamine and 10 mM glucose. Oxygen consumption rate (OCR) and extracellular acidification rate (ECAR) were measured using the XF^e^96 Extracellular Flux Analyzer (Seahorse Bioscience) according to the manufacturer’s instructions.

### Preparation and treatment of precision cut liver slices

Precision cut liver slices (200 μm thickness) were prepared from both normal and tumor - bearing murine livers based on a published protocol (de Graaf et al., 2010) with slight modifications. Mouse livers were perfused with ice - cold University of Wisconsin organ preservation solution (UW) before removal, submerged in UW and kept on ice. Cylindrical cores with a diameter of 5 mm were prepared using a manual biopsy punch (Kai Medical Europe) and placed in a Krumdieck tissue slicer (model MD6000, Alabama research and development) containing ice - cold oxygenated Krebs - Henseleit buffer (KHB, Sigma - Aldrich). William’s E Medium (WME, ThermoFisher Scientific) supplemented with 2.75 mg / ml D - glucose and 50 μg / ml gentamycin (Sigma - Aldrich) was used as the standard culture medium. To assess urea synthesis under stimulated conditions, DMEM with additional urea cycle substrates (2 μM ornithine [Sigma - Aldrich] and 10 μM NH_4_Cl [Sigma - Aldrich]) was used. To determine the effect of complex I inhibition, PCLS were cultured in DMEM with or without 0.5 mM metformin. Slices were incubated in a 6 - well culture plate containing 3 slices per well and 3.5 ml medium. After 1 hour of pre - incubation, slices were transferred to fresh medium and incubated for 24 hours. Culture medium was collected and stored at - 80°C for later analysis of urea production or LDH activity. Viability of the cultured tissue was confirmed by analysis of ATP and total protein content.

### Urea quantification

For the analysis of urea synthesis, 500.000 ASV - B cells or primary hepatocytes were seeded in 6 - well plates with 2 ml of growth medium (DMEM, 10% FCS and 1% penicillin / streptomycin). The medium was replaced 18h later and the cells were incubated for further 24h. 1.5 ml supernatant was collected and centrifuged at 1.000 x g for 5 minutes to separate cells and debris. The remaining supernatant (1.2 ml) was transferred to a new tube and the samples were stored at - 80°C until further use. Urea measurements were conducted by the University Hospital RWTH Aachen Central laboratories applying standard diagnostic procedures.

### Intracellular triglyceride measurement

Intracellular triglycerides in ASV - B cells and primary hepatocytes were determined using the Triglycerides liquicolor^mono^ assay (HUMAN Diagnostics, Wiesbaden, Germany). Briefly, 4x10^5^ cells/well were seeded in 1 ml DMEM medium with 10% FCS and 1% penicillin/streptomycin in 6 well plates and left for 4 hours to attach. Subsequently, the medium was replaced by DMEM supplemented with 0.5% FBS for 16 hours to starve the cells. Then the cells were either left untreated or stimulated with 300 μM oleic acid - albumin (Sigma - Aldrich) for 24 hours. The next day, the cells were washed with PBS once and harvested in 100 μl homogenization buffer (10 mM Tris, 2 mM EDTA, 250 mM sucrose at pH 7.5). Homogenization was conducted by sonification (6 impulses with 10% intensity by sonification, Sonoplus Type UW3100, Bandelin electronic). Cell debris was separated by centrifugation. The assay was conducted according to the manufacturer’s instructions.

### Histochemistry

PAS staining was performed according to routine histochemistry protocols.

### Transmission electron microscopy (TEM)

TEM was performed by the Core Facility for Electron Microscopy of the Charité Berlin as outlined in detail before (Theilig et al., 2001).

### Data analysis

Proteomics raw data were analyzed using the MaxQuant proteomics pipeline v1.4.1.2 and the built in the Andromeda search engine (Cox and Mann, 2008, 2011) with the mouse Uniprot database. Carbamidomethylation of cysteines was chosen as fixed modification, oxidation of methionine and acetylation of N - terminus were chosen as variable modifications. Two missed cleavage sites were allowed and peptide tolerance was set to 7 ppm. The search engine peptide assignments were filtered at 1% FDR at both the peptide and protein level. The ‘match between runs’ feature was not enabled, ‘second peptide’ feature was enabled, while other parameters were left as default. Before comprehensive data analysis, data quality was evaluated using the in - house developed quality control software PTXQC (Bielow et al., 2016).

### Gene set analysis pipeline

In total, 28 proteomic samples were analyzed, coming for 14 mouse livers, with 7 healthy and 7 tumorous livers. Hence, the dataset comprised 7 biological replicates for each condition, with 2 technical replicates per mice liver. For each protein, technical replicates were averaged, while ignoring missing values. Box and density plots were generated to assess the homogeneity of the sample distributions. Clustering was performed over complete cases of proteomics samples using the complete method and Euclidean distance (see pheatmap and hclust R packages) as well as principal component analysis. Differential analysis was performed using the Limma R package (Ritchie et al., 2015). This package assesses the significance of fold changes using parallel linear models sharing variance parameters. This method was originally developed for micro array data but turns out to be particularly suited for shotgun proteomics as it alleviates the scarcity of the measurement matrix by sharing the variance between proteins. Out of 4415 detected proteins, 1886 were tested for significance of their fold changes. 1018 proteins were found to have significant fold changes (FDR 0,05). The list of reviewed proteins associated with metabolic pathways was obtained from Uniprot DB using the gene ontology annotation “metabolic process”. Metabolically relevant pathways for mouse were obtained from the GSKB R package (Bares, 2015). The Piano package (Varemo et al., 2013) was used to estimate the significance of the directional regulation of the pathways. The methods used to generate a consensual p - value in piano were: mean, median, sum, maxmean, stouffer, fisher, reporter, tailStrength, wilcoxon and PAGE. The FDR and t - values yielded by the Limma package were used as gene level statistic. The scripts and data can be accessed here: https://github.com/adugourd/thorsten_liver_model.

### Statistical analysis

All data are presented as mean ± SEM. Statistical analysis was performed by two - tailed Students *t* test using the GraphPad Prism 5.0 software (GraphPad Software). Differences were considered statistically significant at p<0.05.

